# A high-quality telomere-to-telomere LSDV genome assembly

**DOI:** 10.64898/2026.03.24.714000

**Authors:** Caroline Wright, Noemi Polo, Sarwar Azam, Graham Freimanis, G. Taru Sharma, Bryan Charleston, Carrie Batten, Tim Downing, Jonas Albarnaz

## Abstract

Lumpy skin disease virus (LSDV) is an emerging livestock capripoxvirus (CaPV) that continues to cause substantial economic losses across Africa, Asia and Europe. However, important uncertainties remain regarding LSDV genome structure, particularly at the telomeric inverted terminal repeats (ITRs) that are central to host interaction, replication and adaptive evolution. Structural variation at the ITRs is driven by recombination during virus replication. Previous assemblies have relied primarily on short-read sequencing, which provides limited resolution of repeat-rich telomeric regions and propagates structural ambiguities into downstream annotation and comparative analyses. Here, we present a high-quality telomere-to-telomere (T2T) assembly of the LSDV Oman 2009 isolate, generated using a hybrid approach that integrates short Illumina reads with higher-accuracy long Nanopore reads. The resulting 151,091 bp genome contains 157 annotated open reading frames and fully resolves complex repeat-rich structures at both ITRs. This new reference corrects misassemblies present in earlier LSDV genomes and confirms clade-specific gene truncations. This genome provides a robust foundation for improved genomic surveillance, accurate read mapping, mutation detection and evolutionary inference of LSDV, and demonstrates the value of long-read approaches for resolving complex CaPV genome structures.

## Introduction

Lumpy skin disease virus (LSDV, *Capripoxvirus lumpypox*) is an endemic livestock disease in Africa and an emerging threat to bovine health, predominantly in Asia and southern Europe. Owing to its substantial economic impact and rapid transboundary spread, lumpy skin disease is a notifiable disease by the World Organization for Animal Health (WOAH) (WOAH 2010). Although genomic variation clearly influences LSDV virulence and transmission dynamics, large gaps remain in our understanding of its genome structure, particularly within the highly repetitive terminal regions. These limitations can now be addressed using new technological developments in long-read sequencing approaches.

LSDV is a dsDNA virus that belongs to the *Capripoxvirus* (CaPV) genus of the *Poxviridae* family and is typically classified into three main clades (1.1, 1.2 and 2), with clade 2 representing a recombinant lineage derived from ancestors of clades 1.1 and 1.2 (Biswas et al 2021, Haga et al 2024). Currently circulating field strains are predominantly associated with clade 1.1, whereas clade 1.2 is linked to live attenuated vaccine strains. Most existing LSDV genomes are assembled solely from short-read data, leading to fragmented contigs, incomplete annotation, and uncertainty at key loci with high structural complexity. This is especially true for the genes at the terminal 5’ and 3’ ends, e.g. LSDV001 and LSDV156, which are important for host immune modulation (Zhang et al 2025). Short-read assemblies may obscure true biological variation in these regions and introduce artefacts that confound phylogenetic interpretation.

The linear 151 kb LSDV genome has a structure typical to poxviruses. It contains 156 annotated ORFs and is flanked by homologous inverted terminal repeats (ITRs) that harbour four ORFs each (ORFs 1-4 and 153-156, respectively) (Tulman et al 2001) and contain regulatory sequences essential during virus replication. Poxvirus genomes have the two DNA strands covalently linked at the termini forming an incompletely base-paired hairpin analogous to telomeres at the end of each ITR. Poxviruses replicate their genomes through head-to-head and tail-to-tail concatemeric intermediates, and the conserved sequence that is necessary for concatemer resolution is localised between the hairpins and a series of short tandem repeats (Moyer & Graves 1981, Moss 2013). The ITRs and the adjacent 5’ and 3’ regions spanning ORFs 5-23 and 124-152 are among the most variable and encode functions associated with virulence and immune modulation, whereas the central region is highly conserved and encodes genes involved in transcription, DNA replication, and virion assembly. Despite their biological importance, the structure of LSDV ITRs remains poorly defined. By contrast, the ITRs of other poxviruses, such as vaccinia virus (Moss 2013), have been resolved in detail, including long A/T-rich repeats, suggesting a potentially analogous but uncharacterised architecture in LSDV.

Here, we generate a telomere-to-telomere (T2T) annotated reference genome for the Oman 2009 isolate of LSDV by integrating short Illumina reads with long Oxford Nanopore Technology (ONT) reads. This approach used these long reads as the main scaffold for the resulting assembly. It provided an enhanced resolution of the genome, particularly at the repetitive terminal regions, enabling the resolution of complex ITR structures that were previously inaccessible. The resulting high-quality genome provides an improved reference for accurate read mapping, comparative genomics, and evolutionary analyses of LSDV, and as a template for future CaPV hybrid assemblies.

## Methods

### Sample collection

The LSDV/OMAN/2009/MDBKP2 isolate (‘Oman 2009’) was obtained from the Non-Vesicular Reference Laboratory (NVRL) at The Pirbright Institute. The virus was isolated from a cattle skin nodule biopsy after passaging twice in Madin-Darby bovine kidney (MDBK) cells confirmed free of bovine viral diarrhoeal virus (BVDV) by qRT-PCR, following WOAH protocol guidelines (WOAH Terrestrial Manual 2024).

### Read library preparation and sequencing

For short-read Illumina sequencing, total DNA was extracted from 100 µL of cell culture lysate using a MagMax CORE nucleic acid purification kit and a KingFisher Flex automated extraction robot (Thermo Fisher Scientific) according to the manufacturer’s instructions. The extracted DNA was tagmented and indexed using an Illumina DNA Prep with Enrichment kit (Illumina), with 11 cycles of amplification used during DNA tagmentation. The indexed library underwent targeted enrichment with a custom myBaits panel (Daicel Arbor Biosciences), also used in Breman et al 2026 (sequences available as supplementary material). Manufacturer’s instructions were followed for the High Sensitivity Protocol, which included two rounds of probe hybridisation at 63°C. The final library size distribution was determined using a D1000 High Sensitivity ScreenTape on a TapeStation 4200 (Agilent) and quantitated using the Qubit HS kit (Thermo Fisher Scientific). The library was then diluted and re-quantified using the NEBNext Library Quant Kit for Illumina (New England Biolabs) assay. The diluted library was loaded onto an Illumina Nextseq 2000 at a final concentration of 1.6 pM for paired-end sequencing (150 cycles) with a 2% PhiX spike-in.

For long-read ONT sequencing, library preparation was performed using the ONT ligation by sequencing kit (SQK-LSK114) and an input of 340 ng genomic DNA. Final library size distribution was determined using the D1000 genomic DNA ScreenTape on a TapeStation 4200 (Agilent) and quantified using the Qubit DNA BR kit (Thermo Fisher Scientific). Thirty femtomoles of the library were loaded onto a MinION R10.4.1 (FLO-MIN114) flow cell and sequenced for 42 hours on a MinION Mk1B device. Basecalling was performed using Dorado 7.3.11.

### Read library quality control and host read removal

Adapter trimming, removal of both short and long reads with low base quality scores (phred score <30), exclusion of ambiguous (N) bases, mismatched base pair correction in overlapping regions, and cutting of poly-G tracts at 3′ ends was completed using Fastp v0.23.4 (Chen et al 2018). Low-quality ONT bases were removed with Nanoq including trimming the 5’ 11 and 3’ 12 bases from all reads (Steinig & Coin 2022). Low-quality Illumina bases with base qualities <30 and reads with lengths <110 bp were removed using the FASTX-Toolkit v0.0.13 (http://hannonlab.cshl.edu/fastx_toolkit/). FastQC v0.12.1 (www.bioinformatics.babraham.ac.uk/projects/fastqc/) was used to verify the effectiveness of Fastp, the FASTX-Toolkit and Nanoq. We estimated species-level abundances for each read using the k-mer classifier Kraken2 (Wood et al 2019).

The initial ONT read library lengths ranged 35-82,438 bp (median 557 bp) (Table 1). A total of 209,203 LSDV reads had lengths 35-80,952 bp (median 533 bp). This corresponded to 1.9% of the post-QC reads (or 187.06 Mbp of data), reflecting the absence of ONT library target enrichment. This generated a mean genome-wide read depth of 953-fold when mapped to the final assembly with a mean base quality (BQ) of 34.3. The paired-end Illumina data was 99% LSDV because probes were used to enrich that library for LSDV. This resulted in 39.3 million 151 bp LSDV reads, resulting in a mean genome-wide read depth of 10,627-fold with a mean BQ of 39.3.

**Table 1.**
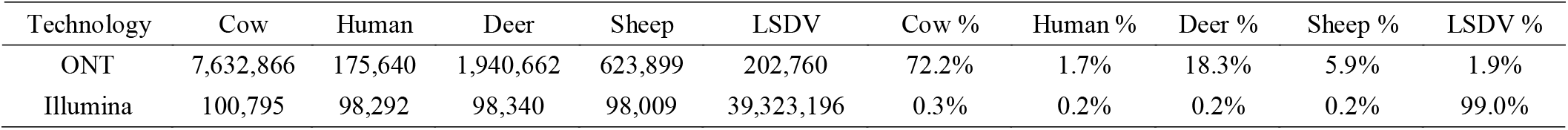
The numbers and percentages of reads classified across organisms by Kraken2 for our ONT and Illumina read libraries. The Illumina reads were enriched for LSDV prior to sequencing.

**Table 2.**
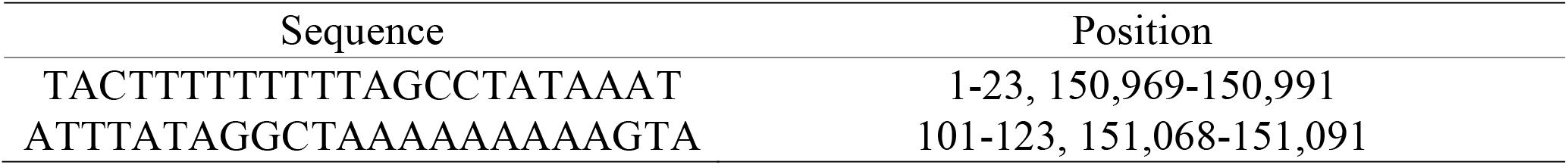
CRS nucleotide sequence at the ITRs in the LSDV Oman 2009 genome assembly.

### *De novo* genome assembly and evaluation

Flye v2.9.1-b1784 was used to assemble the genome from the ONT reads (Kolmogorov et al 2019). This used a subset of the longest reads for initial disjointig assembly, ten polishing iterations, an expected genome size of 151 kb, and high-quality ONT reads. No Flye polishing improvements were observed beyond four iterations. We created assemblies with ONT reads subject to a size requirement of >500 bp, >1,000 bp, >2,000 bp and >4,000 bp that either had no effect or assembled less well. Additionally, reducing the coverage for the initial disjointig assembly was attempted but did not improve assembly quality. Assemblies with alternative contigs were excessively long and impaired overall quality. SPAdes v3.13.1 (Prjibelski et al 2020) was run in careful mode using a range of input: ONT reads, Illumina reads, both read sets, and the same combinations again using contigs from the optimal Flye assembly. We also used Unicycler v0.4.8 (Wick et al 2017) to create Illumina-only, ONT-only and hybrid assemblies using conservative, normal and bold modes.

Assembly quality was measured with Quast based on the assembly length, number of contigs, largest contig size, GC content, N50 and number of N bases. To explore local assembly quality, read pileups and breakpoints, we used Minimap2 v2.24-r1122 (Li et al 2018) to index the assembly and map the reads to it. The resulting Sequence Alignment Map (SAM) files were processed with SAMtools v1.19.2 (Li et al 2009) to process, sort and estimate read depth per site. The Binary Alignment Map (BAM) files stemming from this were examined with IGV v2.16.0 (Robinson et al 2011) for read alignment and paired-end read consistency inspection. Read-depth variation was visualised using R v4.3.2 (R Core Team 2025) packages ggplot2 v3.5.1 (Wickham 2016) and gridExtra v2.3 (Auguie & Antonov 2017).

### Post assembly polishing and genome annotation

The assembly was polished using four steps with Medaka, Pilon, Polypolish and Pypolca. Firstly, Medaka v2.0.1 (https://github.com/nanoporetech/medaka) was used to polish the optimal Flye assembly to correct large indel and homopolymer errors using the cleaned ONT reads with a r1041_e82_400bps_sup_v5.0.0 model. The assemblies were compared using Nucmer v3.1 (Marçais et al 2018). Secondly, we used Pilon v1.24 (Walker et al 2014) to fix SNPs, indels, gaps, local misassemblies, and ambiguous bases over five iterations using the assembly resulting from Medaka. Thirdly, Polypolish v0.6.0 corrected errors in repeats in the Pilon output (Wick & Holt 2022). To run Polypolish, BWA v0.7.17 mapped the Illumina reads to all possible matches where forward and reverse reads were mapped separately. These reads were filtered by insert size and used for polishing. Fourthly, Pypolca v0.3.1 in careful mode was used to do a final correction step on the Polypolish assembly using the Illumina reads (Bouras et al 2024). Canu (Koren et al 2017) was used to error-correct and assemble the ONT reads but this provided no improvement.

The assembly versions were annotated using Prokka v1.14.6 (Seeman 2014) with the --kingdom Viruses option specified, with gene annotation guided by the reference genome KX894508 (155920/2012; 156 ORFs). Annotation was also mapped from the reference strain LSDV NI-2490 (AF325528.1) using Genome Annotation Transfer Utility (GATU) (Tcherepanov et al 2006). The annotation was manually inspected by visualising long- and short-read alignments in IGV v2.16.0 (Robinson et al 2011) to ensure the gene boundaries were correct. Genome completeness and quality, including ITRs within the terminal regions, was further assessed using CheckV v1.0.3 (Nayfach et al 2021).

## Results and Discussion

### A more accurate LSDV reference genome

We used a combination of long-read ONT and short-read Illumina sequencing data to *de novo* assemble a new reference genome sequence for the LSDV Oman 2009 isolate. The final assembly comprised a single contig of 151,091 bp (N50 151,091 bp), with 157 predicted ORFs and a mean GC content of 25.9% (median 25.6%, Figure S1). This contig was generated from long reads using Flye v2.9.1-b1784 (Kolmogorov et al 2019) because Flye has a low chance of making large-scale errors (Wick et al 2023). It was refined through iterative polishing with Medaka v2.0.1 that made 15 changes, Pilon v1.24 (Walker et al 2014) that fixed two SNP errors, and correction with Polypolish v0.6.0 (Bouras et al 2024) that fixed nine indel and two substitution errors. Alternative assembly strategies resulted in fragmented assemblies with inflated total lengths and numbers of contigs. Overall, polishing with Illumina reads corrected only 28 bp (0.019%) of the 151,091 bp genome assembled from long ONT reads.

The genomic termini of the Oman 2009 reference genome were verified using a known 123 bp hairpin sequence (MN072625.1, Biswas et al 2020), which had T-A differences at bases 30 and 33. The terminal 23 bp of the hairpin contained concatemer resolution sequences (CRSs), which mediate site-specific genome resolution during viral replication (Moss 2013). The 5’ and 3’ ITRs were sequence-symmetrical and contained CRSs at homologous positions, matching a previously described GTPV CRS (Biswas et al 2020).

To assess assembly quality, we visualised per-base read depth for both short- and long-read data across the genome (Figure 1). Long-read depth was relatively uniform, with a standard deviation of 66 (7% of the median depth of 962), compared with greater variability in short-read depth (standard deviation 2,433; 23% of the median depth of 10,379). Examination of the genome annotation additionally identified two truncated genes, LSDV019 and LSDV026, which are also truncated in other wild-type LSDV isolates from clade 1.1 (Bamouh et al 2021, Kumar et al 2023, Chang et al 2025). Gene fragments LSDV019a/b and LSDV026a/bwere present as truncated coding sequences matching LSDV019 and LSDV026, respectively, in the Neethling isolate NI-2490 from clade 1.2 (AF325528.1), where the genes are full length. LSDV019 is homologous to kelch-like virulence factors characterised in orthopoxviruses (Pires de Miranda et al 2003, Beard et al 2006, Froggatt et al 2007). LSDV026 is also truncated in SPPV and GTPV and is an orthologue of vaccinia virus F11, a virulence factor involved in regulation of virus spread (Cordeiro et al 2009). Furthermore, we annotated a new ORF, tentatively named LSDV042.5, which is predicted to be an orthologue of vaccinia virus O3L. O3L encodes a 35-amino-acid protein that is a component of the entry fusion complex required for virus penetration into host cells and conserved across poxviruses of vertebrates (Satheshkumar & Moss 2009). LSDV042.5 encodes a predicted polypeptide of 29 amino acid residues and therefore, was not annotated in the first LSDV genome sequenced (AF325528.1) even when a sensitive cutoff (ORFs >30 amino acid residues) was applied (Tulman et al 2001). Nonetheless, LSDV042.5 is likely to be functional because the O3L-like gene from SPPV (AY077832.1), which is 93% identical to LSDV042.5, can complement O3L deficiency in a recombinant vaccinia virus (Satheshkumar & Moss 2009).

**Figure 1.**
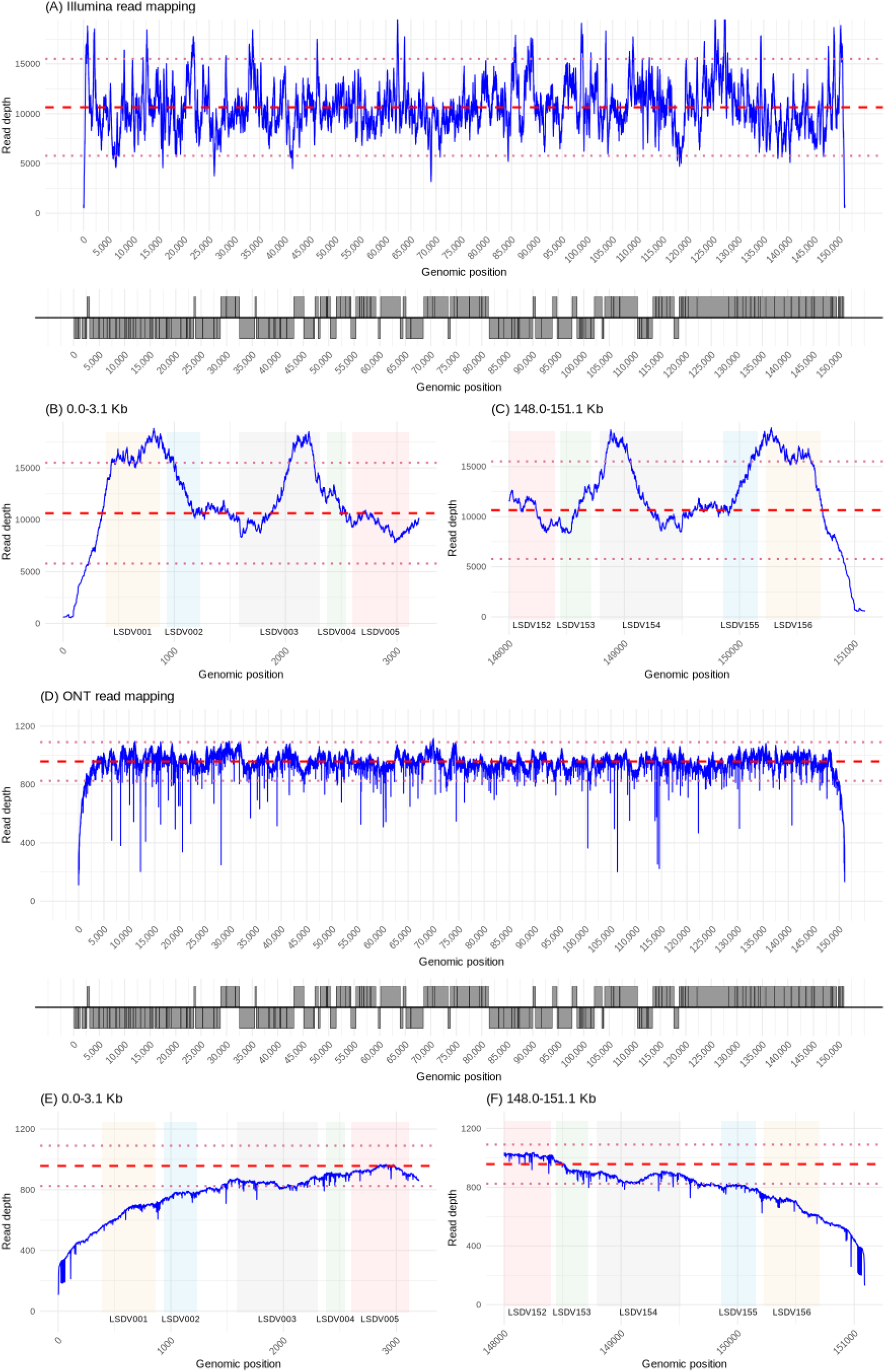
Per-base read depth across the LSDV Oman 2009 assembly for Illumina data (A) genome-wide, (B) at 0-3.1 kb and (C) at 148-151.1 kb, and equivalent ONT data within the same regions (D) genome-wide, (E) at 0-3.1 kb and (F) at 148-151.1 kb. The blue lines show read depth per site. The red dashed lines show genome-wide average read depth. The red dotted lines indicate twice the SD above and below the average. The small black boxes in (E) indicates the CRS. The ITRs are at 1-2,547 bp and 148,557-151,091 bp. The grey panels below (A) and (D) represent locations of annotated CDSs across the genome. Coloured shading in (B), (C), (E) and (F) demarcates LSDV001 (beige), LSDV002 (pale blue), LSDV003 (pale purple), LSDV004 (pale green), LSDV005 (pink), LSDV152 (pink), LSDV153 (pale green), LSDV154 (pale purple), LSDV155 (pale blue) and LSDV156 (beige).

The reference genome contained a 5’ ITR spanning positions 1-2,547 bp and a corresponding 3’ ITR spanning positions 148,557-151,091 bp, yielding ITR lengths of approximately 2.5 kb at each terminus. This annotation was supported by the genomic positions of conserved flanking genes, including LSDV005 (2,599-3,111 bp) at the 5’ end and an ankyrin repeat protein (LSDV152; 146,929-148,398) at the 3’ end. Analysis of read depth revealed abrupt changes in short-read coverage at approximately 1.2, 3.3, 5.8 and 6.4 kb from the 5’ end, with corresponding features observed at the 3’ end of the genome (Figure 1). These coverage transitions were present when short reads, but not long reads, were mapped to the reference. Median short-read depth across the 5’and 3’ ITRs (12,580 and 12,592, respectively) exceeded the genome-wide median short-read depth of 10,379, indicating overrepresentation of these regions due to their repetitive structure. This suggests that short reads alone overestimate ITR length by 22%, whereas long reads spanned the repetitive regions and enabled accurate estimation of ITR structure and length.

Alignment of the Oman 2009 hybrid assembly with the PacBio-based genome assembly PX492334 (Breman et al 2026) revealed only 12 mismatches across the full genome length. This high level of concordance indicates that both ONT and PacBio sequencing approaches can generate highly accurate assemblies for LSDV. Comparison of the terminal regions showed that the first 67 bp of the Oman 2009 assembly, including the first CRS, were not present in the 5’ ITR of PX492334, where only a partial 11 bp match was observed; an identical pattern was detected at the 3’ end. Importantly, the presence of this additional sequence symmetrically at both genome termini in the Oman 2009 assembly supports its authenticity and suggests that the hybrid ONT-Illumina approach resolved an additional 56 bp at each end of the genome relative to the PacBio-only assembly.

### Summary

This study used a hybrid short- and long-read sequencing strategy to generate an annotated reference genome for the LSDV Oman 2009 isolate, resolving the architecture of the ITRs more accurately than previously available assemblies. The results provide a framework for future hybrid genome assemblies of CaPVs and demonstrate the effectiveness of ONT R10 long-read sequencing for producing high-quality CaPV genomes. Consistent with observations in bacterial systems (Sereika et al 2022), our findings suggest that long-read data alone may be sufficient for accurate CaPV genome assembly, while short-read data are more effectively applied to resequencing and read mapping (Wright et al 2025). The improved resolution of ITRs and truncated genes in the Oman 2009 genome supports its use as a reliable LSDV reference and facilitates more accurate clade-specific analyses and diagnostic assay development (Tennakoon et al 2025). Continued improvements in sequencing technologies, including ultra-long reads, alterative long read platforms and improved polymerase accuracy, are likely to enable further refinement of LSDV and CaPV genome assemblies.

## Data Availability

The Illumina and ONT reads are on the SRA under the BioProject accession PRJNA1366067. The ONT reads are available under the accession SRX31142208. The Illumina reads are available under the accession SRX31141470. The genome assembly file and its annotation files are on the Nucleotide database under the accession PV877838. The code used to produce these analyses is publicly available at https://github.com/downingtim/LSDV_hybrid_assembly/

### Acknowledgements

We would like to acknowledge the Pirbright Institute’s Bioinformatics, and High-Throughput Sequencing STPs and Non-Vesicular Reference Laboratory (NVRL). We also acknowledge the use of generative artificial intelligence tools (ChatGPT, Copilot) to write code used for this work, and to proofread the text. All outputs were reviewed, tested, and/or validated by the authors.

---

**Figure S1.**
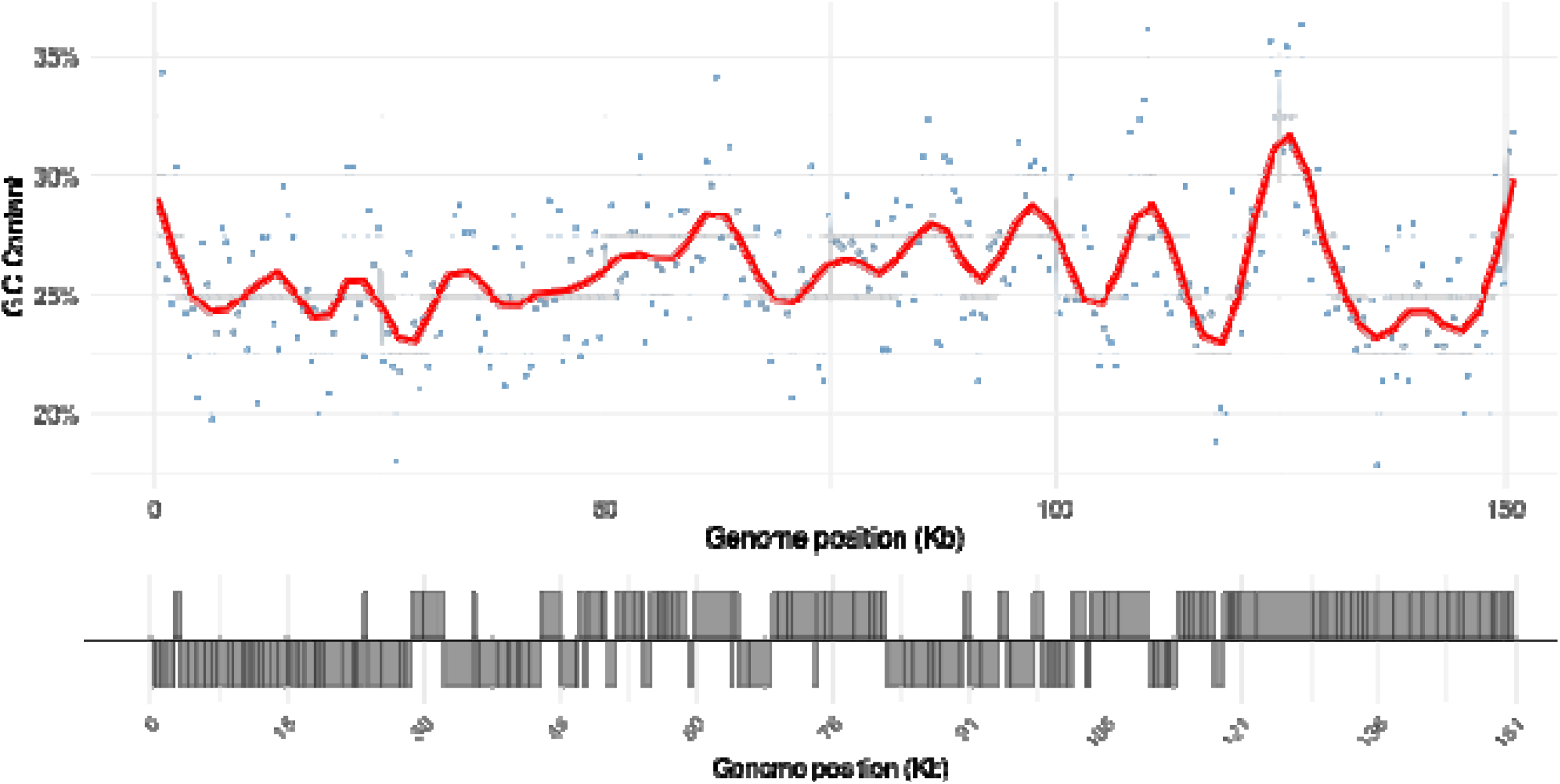
The genome-wide variation in GC content in the LSDV Oman 2009 hybrid genome assembly. The red line shows the smooth pattern in 500 bp bins and the grey area shows the 95% confidence interval. The grey panel represents the locations of the annotated CDSs across the genome. The y-axis has been cut.

